# Rag2Mol: Structure-based drug design based on Retrieval Augmented Generation

**DOI:** 10.1101/2024.10.20.619266

**Authors:** Peidong Zhang, Xingang Peng, Rong Han, Ting Chen, Jianzhu Ma

**Affiliations:** Department of Computer Science and Technology, Tsinghua University, Beijing, China; Institute of Artificial Intelligence, Tsinghua University, Beijing, China; Institute for Artificial Intelligence, Peking University, Beijing, China; School of Intelligence Science and Technology, Peking University, Beijing, China; Department of Electronic Engineering, Tsinghua University, Beijing, China; Institute for AI Industry Research, Tsinghua University, Beijing, China

## Abstract

Artificial intelligence (AI) has brought tremendous progress to drug discovery, yet identifying hit and lead compounds with optimal physicochemical and pharmacological properties remains a significant challenge. Structure-based drug design (SBDD) has emerged as a promising paradigm, but the inherent data biases and ignorance of synthetic accessibility render SBDD models disconnected from practical drug discovery. In this work, we explore two methodologies, Rag2Mol-G and Rag2Mol-R, both based on retrieval-augmented generation (RAG) to design small molecules to fit a 3D pocket. These two methods involve searching for similar small molecules that are purchasable in the database based on the generated ones, or creating new molecules from those in the database that can fit into a 3D pocket. Experimental results demonstrate that Rag2Mol methods consistently produce drug candidates with superior binding affinities and drug-likeness. We find that Rag2Mol-R provides a broader coverage of the chemical landscapes and more precise targeting capability than advanced virtual screening models. Notably, both workflows identified promising inhibitors for the challenging target PTPN2, which was considered undruggable because of structural characteristics of phosphatases. Our highly extensible framework can integrate diverse SBDD methods, marking a significant advancement in AI-driven SBDD. The codes are available at: https://github.com/CQ-zhang-2016/Rag2Mol.

## 1 Introduction

Artificial intelligence (AI) has achieved tremendous success in drug discovery, providing profound and inspiring insights across various stages^1, 2^. Of these, finding hit/lead compounds, as the foundation for starting a new drug discovery project, remains a significant challenge in biopharmaceutical research, particularly in acquiring favorable physicochemical and pharmacological properties^3^. Researchers have explored various approaches to discover protein pocket-specific molecules^4–6^, and structure-based drug design (SBDD) has emerged as a promising paradigm due to its superior performance among these efforts^7^. Unlike virtual screening methods^8–12^ limited by the chemical diversity of existing compound libraries or ligand-based drug design (LBDD) methods^13–17^ which lack protein structural information, SBDD models explicitly incorporate the three-dimensional geometric information of target proteins, directly generating ligands with appropriate topological structures and high binding affinity within the protein pocket^18–23^.

Recently, several representative methods have been proposed for SBDD tasks, demonstrating promising results. Work^24^ is the first approach to generate molecules based on 3D structures with favorable biological properties. It iteratively adds atoms or bonds to construct 3D molecules, making autoregressive models^24–28^ a key category in the SBDD field. Diffusion-based methods^29–32^ sample the distribution of atoms from the prior noise distribution and generate entire molecules in iterative feed-forward processes followed by post-processing to assign bonds. Additionally, flow-based^33^ and language model-based^34^ approaches encode contextual information to obtain rotation-invariant features, offering different perspectives in molecular generation.

However, several challenges have rendered SBDD models disconnected from practical drug discovery. Firstly, SBDD models inherently ignore the synthetic accessibility, resulting in generated structures usually falling outside the synthesizable chemical spaces^33, 35^. Secondly, as data-driven methods, SBDD models are intrinsically limited by the quality and coverage of their training data^36^, which is undoubtedly a drop in the ocean compared to the potential space of complexes. Finally, pocket-specific molecules designed by SBDD models require extensive downstream validation^37^, with cumbersome procedures and accumulated uncertainties deterring drug developers. Recent studies have proposed various innovative approaches to partially address these challenges, including introducing chemical knowledge^36, 38^ and relevant features^39^, projection into synthesizable chemical space^35^, and extracting interaction patterns from retrieved molecules^32^. While these pioneering efforts are promising, much room for improvement remains.

In this work, we have designed two retrieval augmented generation^40^ (RAG)-based workflows to enhance the performance of SBDD models in real-world applications (as illustrated in Figure 1). In Rag2Mol-G, we use a pre-trained two-level retriever during both training and sampling. The global retriever is to retrieve database for small molecules that potentially bind to the pocket, and the molecular retriever is to rank and choose them as context information. The advantage of this workflow is that the retrieved molecules have both potential interaction affinity and synthetic accessibility, implicitly assisting the AI model in learning structural knowledge and topological rules. In Rag2Mol-R, we first generate candidate molecules based on Rag2Mol and then search for similar molecules in the public database. Such methodology is similar to the pipeline PocketCrafter developed by Novartis^41^, which successfully designed three WDAC inhibitors and found the success rate of detecting similar lead compounds significantly surpasses that of direct virtual screening. In conventional virtual screening, at least three AI models need to be trained for screening, docking, and binding affinity prediction based on docked structures. The first and second type errors generated by each model will accumulate, ultimately leading to a high rate of false positives. The intuition behind the Rag2Mol-R methodology is that it consolidates multiple AI models into one model, greatly improving the success rates. Rag2Mol-G is particularly useful when novel drug candidates with high affinity and drug-likeness are required, leveraging the generative power of our model. On the other hand, Rag2Mol-R excels in identifying synthesizable analogs that can be readily purchased or synthesized, providing a practical approach for drug repurposing and optimization.

**Figure 1.**
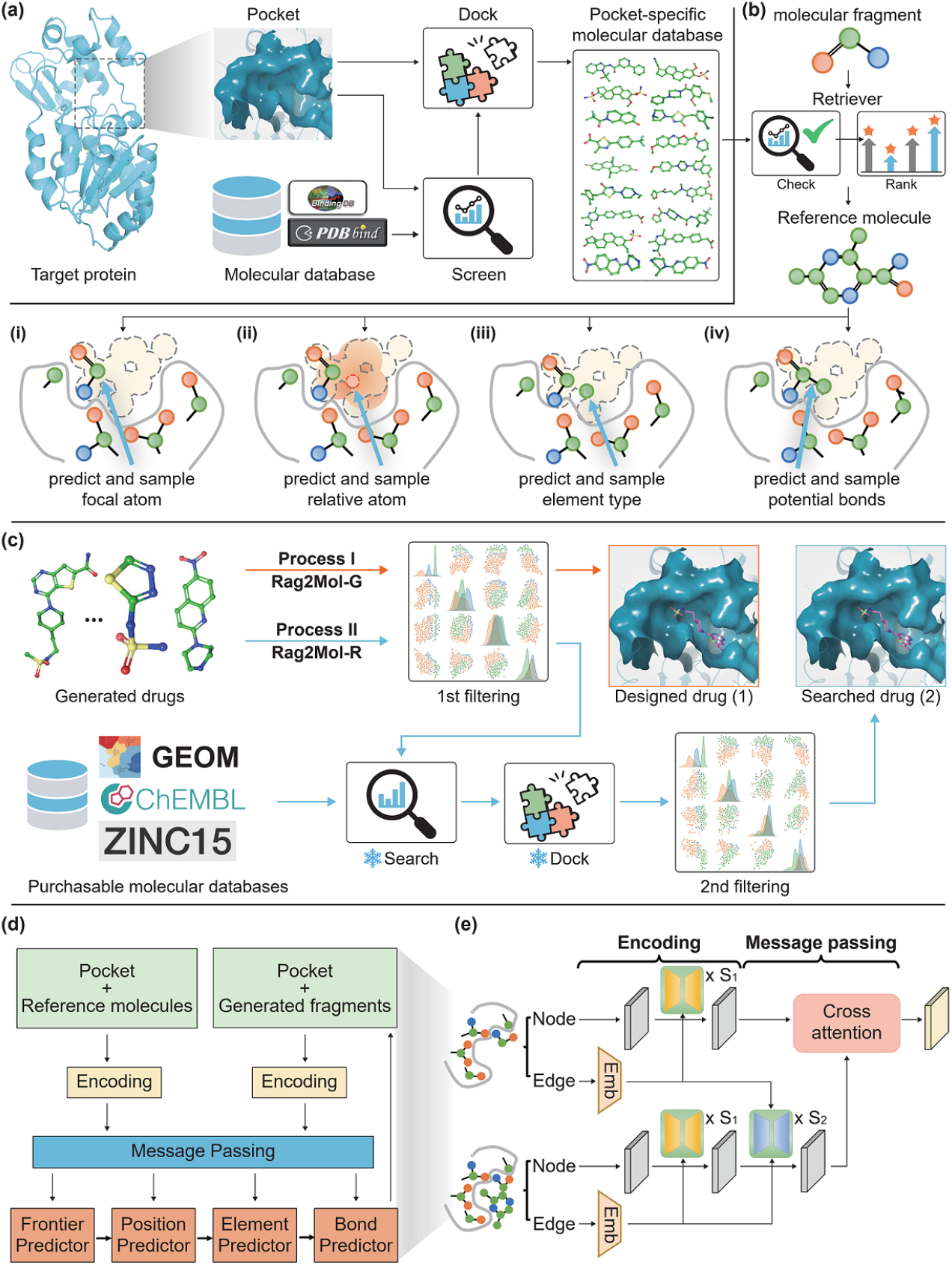
The Rag2Mol pipeline. (a) **Global retriever.**Constructing pocket-specific molecular database for each given target protein. (b) **One step of the Rag2Mol**. Using the reference molecule selected by the molecular retriever, the autoregressive model sequentially predicts various information for the next atom (including focal atom, relative position, element type, and the valence bonds) based on the generated molecular fragment. (c) **Two workflows for applying Rag2Mol**. In the Rag2Mol-G workflow, we filter drug candidates following the widely accepted threshold settings. In the Rag2Mol-R workflow, filtered molecules are then subjected to clustering and sampling. These sampled scaffolds are employed for similarity searches within synthesizable compounds. (d) **Simplified architecture of Rag2Mol**. (e) **Information transfer mechanism in hidden space**.

Extensive experiments provided by this study demonstrate the superiority of our workflows. The vanilla Rag2Mol achieves state-of-the-art (SOTA) performance across multiple evaluation metrics on widely-used dataset, surpassing other advanced SBDD models while maintaining interpretability; Molecules screened by Rag2Mol-R significantly outperform those selected by other advanced virtual screening tools, covering a broader chemical landscape. Thus, Rag2Mol-G fits targets with multiple binding templates, while Rag2Mol-R excels with traditionally “undruggable” targets. Furthermore, both workflows could identify promising drug candidates for the challenging case PTPN2, outperforming current active site inhibitors. Finally, although we employ widely recognized autoregressive-based methods, almost all AI-based SBDD methods could be integrated into our framework. Meanwhile, each module of the protocol exhibits convenient extensibility.

## 2 Methods

### 2.1 Overview of Rag2Mol

We choose RAG architecture to incorporate prior chemical knowledge and topological rules into the generation process, addressing the limitations of traditional SBDD models. Therefore, Rag2Mol is a RAG-based SBDD generative model, consisting of two components: an augmented molecular generator and two-level retrievers (a global retriever and a molecular retriever). As an auto-regressive model, the generator depicts the molecule generation as generating atoms one by one conditioned on the protein pocket and previously generated fragments. The retriever augments the generation process in the following ways, which distinguishes Rag2Mol from other auto-regressive SBDD models. At the beginning of the generation, we use a global retriever to build a pocket-specific molecular database (Figure 1a). This retriever contains a virtual screening model and a docking model to screen and dock potential small molecules from an external molecular database to the given pocket. In the generation step, a molecular retriever is proposed to retrieve the pocket-specific database based on the previous fragment and select reference molecules to assist the generator (Figure 1b), and the generator produces a new atom by sequentially predicting the focal atoms among existing atoms, predicting the new atom’s relative position, and determining the new atom’s element type and the valence bonds. Based on the Rag2Mol-generated molecules, two application workflows are developed to fit different scenarios (Figure 1c). Finally, Figures 1d and 1e present a simplified framework of Rag2Mol and the detailed information flow within the module.

Formally, let *P* be the pocket and *M*^*t*^ be the generated molecular fragment at the *t*-th generation step (*t* = 1, 2, …, *T* ). In the beginning, the pocket-specific molecular database, denoted as 𝒟, is derived as:

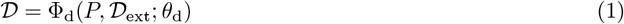

where Φ_d_ is the global retriever with parameters *θ*_d_, and 𝒟_ext_ is the external molecular database. The generation process is defined as:

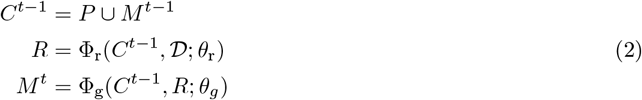

where Φ_r_ and Φ_g_ are the molecular retriever and the generator with parameters *θ*_d_ and *θ*_g_, respectively. *R* is the reference molecule retrieved from the pocket-specific molecule database.

### 2.2 Implementation of the two-level retriever

Inspired by previous works, the global retriever Φ_d_ directly utilizes ConPLex^9^ and FABind^42^ for docking and screening. However, for the molecular retriever, we trained a light network for faster retrieving due to its frequent usage. The molecular retriever essentially needs the capability to measure the similarity between the reference molecule and the final molecules that will be generated from the current intermediate fragment. To train this retriever, we sampled a protein-molecule pair (*P, M* ) from the dataset and randomly masked several atoms of the molecule, denoted as *M*_mask_. We then selected reference molecules *M*_ref_ from its pocket-specific molecule database. The molecular retriever took as input the protein *P*, the masked molecule *M*_mask_, and the reference molecule *M*_ref_ and was trained to predict the similarity between the reference molecule *M*_ref_ and the ground truth molecule *M* .

In the implementation, ProtBert^43^ and Morgan fingerprints^44^ were used to encode the proteins and the small molecules into vectors with dimensions of 1024 and 2048 (which were recommended by original works), respectively. Then multi-layer perceptrons were utilized to process these features and finally predicted the similarity scores. Here, the similarity was defined as the cosine similarity of the molecular fingerprints, and the mean squared error was used as the loss function. Molecular retriever was quickly trained to convergence, and the mean squared error between the predicted similarity scores and the true values on a randomly split test set was approximately 0.12.

### 2.3 Encoders

We modeled the binding pocket and the molecules as the k-nearest neighbor (KNN) graph where heavy atoms are nodes and each atom is connected to its k-nearest atoms with edges. To keep the E(3)-equivariance of the generator, we preserved both scalar and vector features for the node and edge features. The input scalar node features were composed of the element type, the amino acid type, the backbone/side-chain identifier of protein atoms, and an identifier to indicate whether the atom belongs to a protein or molecule. The input scalar edge features were composed of the edge lengths, the bond types, and a bool identifier indicating the valence of the bond. The input vector node features were the coordinates of heavy atoms, and vector edge features were the 3D unit directional vector of the edge.

Using the geometric vector perceptrons (GVP^45^) as the basic block, we built two parallel encoders where multiple aggregating layers *G*_*a*_ and updating layers *G*_*u*_ are concatenated and interleaved to learn the local structure representations. We denote the node and edge features as ***v*** and ***e***, respectively. The features of molecular fragment and retrieved reference molecule are encoded separately according to the following formula:

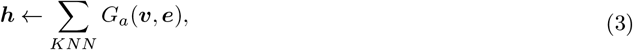

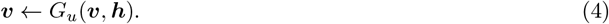

### 2.4 Message passing

We construct a cross-KNN graph to allow messages to flow from the reference molecule to the molecular fragment. Specifically, the reference molecule and molecular fragment are aligned based on the pocket structures. Thus, the cross-KNN graph takes the generated molecular atoms as nodes, and each atom is connected to its k-nearest reference molecular atoms. Figure 1e shows the details of the message-passing module. Let (***v***_*f*_, ***e***_*f*_ ) and (***v***_*r*_, ***e***_*r*_) represent the encoded features of molecular fragment and reference molecule. The edges from the nodes of reference molecules to molecular fragments are represented as ***e***_*r→f*_, which are initialized as the relative distances and the unit directional vector of the atom pairs.

A similar aggregating layer is employed in the cross-KNN graph, and the aggregated node features are used to update information through cross-attention layers, ensuring smoothness and robustness. Using *Attn* to indicate the cross-attention network, the specific formulas are as follows:

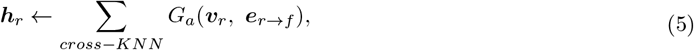

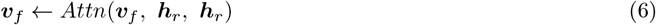

### 2.5 Predictors

The extracted hidden representations capture not only the chemical and geometric attributes within a protein pocket but also the general molecular structural laws and interaction patterns from reference knowledge. These representations are used as the inputs for the predictors including the focal atom predictor, position predictor, and element-and-bond predictor. Following previous works^24, 25, 36, 46^, we use GVP-based networks to predict the probability of focal atoms, parameters of multivariate Gaussian mixture distribution modeling interatomic distance, element types, and probable bonding types. Particularly, we use trigonometry self-attention to capture chemical bonding relationships.

### 2.6 Training procedure

The generator is trained to recover randomly masked atoms. Specifically, for a protein-molecule pair in the dataset, a random ratio of molecular atoms is masked, where the ratio value is sampled from a uniform distribution. The remaining atoms that have chemical bonds to the masked atoms are labeled as focus atoms. The focal predictor is trained to predict focal atoms and recover the masked atoms. We also sample noise positions from the surrounding environment as negative examples, enhancing the learning capability of Rag2Mol. For each masked fragment, we utilize the *Rank* model to retrieve the top 4 reference molecules, and Rag2Mol randomly selects one reference molecule or disregards reference information during each training step. Besides, a teacher-forcing strategy is employed, with each predictor independently trained with ground truth.

Finally, the focus atom predictor and the position predictor use binary cross-entropy loss and negative log-likelihood loss, respectively. Cross-entropy loss is applied to the multi-classification predictions of element types and bonding relationships. Rag2Mol is optimized through the sum loss of predictors. Details about predictors and loss functions are provided in section 3 of Supplementary Information.

### 2.7 Details of downstream workflows

As shown in Figure 1c, we have developed two workflows for drug discovery, named Rag2Mol-G and Rag2Mol-R. In the Rag2Mol-G workflow, we subject the filtered drug candidates to precise binding affinity calculations and subsequent wet-lab experiments. In the Rag2Mol-R workflow, the representative molecule is randomly selected as a scaffold template from each molecular cluster. Based on these templates, we search for similar molecules within existing synthesizable compounds. These molecules are deduplicated and then subjected to accurate docking software Glide^47^, yielding the final set of drug candidates. The detailed implementation, tool explanations, and other set-ups are included in section 4 of Supplementary Information.

## 3 Result and discussion

We evaluate Rag2Mol’s performance on the SBDD tasks and the virtual screen tasks. For the SBDD task, we chose eight SBDD models: Pocket2Mol^46^, ResGen^25^, AR^24^, GraphBP^48^, FLAG^49^, TargetDiff^30^, Decomp-o and Decomp-r^29^, as baselines for subsequent comparison. Fairly, all models were trained and tested on the CrossDock 2020^50^, and the native ligands were also compared. We use widely recognized metrics to assess the common properties: (a) **Vina Dock** and **Vina Score**: Affinity scores before and after docking, respectively. **Affinity**^**1**^ and **Affinity**^**2**^: Percentage of molecules with higher **Vina Dock** and **Vina score** to the existing ligands. (c) **PB-Valid**: Percentage of valid molecules checked by the PoseBusters tool^51^. (d) **CNN affinity**: CNN-based predicted affinity^52^. (e) **Clash**: Number of steric clashes. (f) **SE**: Strain energy. (g) **QED**: Quantitative estimation of drug-likeness. (h) **SA**: Synthetic accessibility. (i) **Lipinski**: Number of obeyed rules of Lipinski’s rule of five. (j) **LogP**: Partition coefficient. (k) **Diversity**: Average molecular similarities for each pocket. Besides, we use Kullback-Leibler (KL) divergence to analyze the distributions of bond angles and dihedral angles and the ratio of rings with different sizes. The root-mean-square deviation (RMSD) is used to measure pose differences before and after reprocessing.

For the virtual screen task, we use the same metrics to compare the common properties of chosen molecules by Rag2Mol-R and three other advanced virtual screen methods: ConPLex^9^, DrugBAN^8^, and UdanDTI^11^. Notably, the structures of target proteins are downloaded from PDB (https://www.rcsb.org) in real-world cases. Details of the datasets, baselines, and evaluation schemes are included in Supplementary Information.

### 3.1 Evaluation of common properties for generated molecules

Table 1 shows that Rag2Mol exhibits optimal or near-optimal results across almost all metrics covering binding energies and drug-likenesses. Docking-related scores evaluate both the binding strength of generated molecules and the SBDD model’s ability to position molecular poses. Some generated molecules with distorted structures (observed in GraphBP) would be reprocessed during docking, resulting in false-positive results. In contrast, Rag2Mol’s top 1/3 molecules consistently demonstrate the lowest binding energies across both affinity evaluations, exceeding the affinity of native ligands by about 3 kcal/mol (around 125-fold activity). Detailed results of top 1/3/5/10 molecules are shown in Supplementary Table S4. Affinity metrics indicate that more than 60% of the Rag2Mol-generated molecules have higher binding affinities than natural ligands, outperforming the closest SBDD method by about 5%. These results indicate that Rag2Mol is more likely to produce tightly binding molecules.

**Table 1.** The mean binding energies and drug-likeness properties of top 1/3/5/10 molecules in drug generation. (↑ )/ (↓) indicates larger / smaller is better. Top 3 results are highlighted with **bold** text, underlined text, and ^*\**^, respectively.

|  | Test set | GraphBP | Pocket2Mol | ResGen | PocketFlow | AR | FLAG | TargetDiff | Devomp-o | Decomp-r | OurModel |
| --- | --- | --- | --- | --- | --- | --- | --- | --- | --- | --- | --- |
| Affinity<sup>1</sup>(↑) | - | 52.8% | 56.8%* | 40.6% | 52.9% | 35.1% | 26.8% | 52.9% | <u>57.8%</u> | 46.1% | **61.5%** |
| Affinity<sup>2</sup>(↑) | - | 39.8% | <u>55.7%</u> | 32.3% | 43.6% | 42.8% | 12.8% | 54.8%* | 54.0% | 40.0% | **60.7%** |
| Diversity(↑) | - | 0.882 | 0.863 | 0.847 | 0.863 | 0.837 | **0.900** | 0.892* | 0.889 | 0.842 | <u>0.893</u> |
| PB-Valid(↑) | 95.0% | 31.2% | <u>73.1%</u> | 21.1% | 50.7% | 55.6% | 18.4% | 50.8% | 48.3% | 71.8%* | **79.7%** |
| CNN affinity(↓) | 5.551 | 5.107 | 3.714 | 3.424 | **2.852** | 3.819 | <u>2.958</u> | 4.571 | 5.364 | 4.778 | 3.036* |
| Clash(↓) | 7.458 | 19.107 | 7.032* | 29.228 | 10.279 | <u>5.923</u> | 69.441 | 13.213 | 16.947 | 10.086 | **5.765** |
| SE-75%(↓) | 61.184 | 152.322 | 49.792* | <u>39.762</u> | 636.445 | 432.008 | 4532.652 | 589.482 | 3582.080 | 13722.683 | **39.060** |
| **Top 1** |  |  |  |  |  |  |  |  |  |  |  |
| Vina Dock(↓) | -7.204 | -9.332 | -9.418 | -9.033 | -9.827 | -8.391 | -8.333 | <u>-10.132</u> | -10.031* | -8.387 | **-10.636** |
| Vina Score(↓) | -5.916 | -1.170 | -8.742 | -7.245 | -8.587 | -8.154 | -6.498 | <u>-9.886</u> | -9.422* | -7.562 | **-10.160** |
| QED(↑) | 0.476 | 0.556 | 0.541 | 0.566* | 0.525 | 0.536 | **0.619** | 0.466 | 0.470 | 0.524 | <u>0.570</u> |
| SA(↑) | 0.728 | 0.468 | 0.740* | <u>0.743</u> | 0.733 | 0.569 | 0.637 | 0.498 | 0.602 | 0.681 | **0.761** |
| Lipinski(↑) | 4.340 | 4.810 | 4.910 | 4.848 | 4.930* | 4.691 | **4.980** | 4.610 | 4.525 | 4.680 | <u>4.940</u> |
| LogP | 0.894 | 1.552 | 2.761 | 2.711 | 3.923 | 0.664 | 2.702 | 2.448 | 3.492 | 2.174 | 3.486 |
| **Top 3** |  |  |  |  |  |  |  |  |  |  |  |
| Vina Dock(↓) | -7.204 | -8.809 | -9.258 | -8.847 | -9.450 | -8.190 | -8.039 | <u>-9.735</u> | -9.695* | -8.272 | **-10.450** |
| Vina Score(↓) | -5.916 | 2.983 | -8.504 | -6.958 | -7.988 | -7.914 | -6.098 | <u>-9.232</u> | -9.035* | -7.400 | **-9.992** |
| QED(↑) | 0.476 | 0.496 | 0.538 | <u>0.572</u> | 0.543 | 0.525 | **0.624** | 0.483 | 0.472 | 0.525 | <u>0.567*</u> |
| SA(↑) | 0.728 | 0.478 | 0.738 | <u>0.755</u> | 0.743* | 0.571 | 0.641 | 0.506 | 0.600 | 0.678 | **0.768** |
| Lipinski(↑) | 4.340 | 4.787 | 4.890 | 4.859 | 4.923* | 4.729 | **4.993** | 4.587 | 4.566 | 4.643 | <u>4.933</u> |
| LogP | 0.894 | 1.470 | 2.656 | 2.718 | 3.873 | 0.649 | 2.438 | 2.481 | 3.433 | 2.073 | 3.399 |

Moreover, SA score, SE, Clash and PB-Valid metrics suggest that Rag2Mol demonstrates a greater ratio of generating valid molecules under equivalent conditions. The QED and Lipinski scores of Rag2Mol-generated molecules are slightly lower than that of FLAG, which can be attributed to the fact that FLAG assembles predefined molecular fragments rather than individual atoms, reflecting a methodological bias. The most detected clashes suggest inherent conformational instability in the FLAG method. Besides, the average LogP values for all generated molecules range between 0.58 and 3.5, within the commonly accepted range. Finally, Rag2Mol also shows high diversity.

### 3.2 Quality of generated conformation

The substructure distribution in generated molecules should align with those in natural molecules, exhibiting appropriate docking poses and conformational stability. As illustrated in Figures 2a-f, Rag2Mol closely mirrors the test set’s distribution of ring sizes, particularly favoring stable 6-membered rings while disfavoring uncommon 3-, 4-, and 7-membered rings, reflecting its effective learning of structural features via retrieval-augmented learning. Diffusion-based methods, however, display lower reliability due to denoising challenges in mixed spaces, while FLAG explicitly avoids larger rings but ignores fused rings (e.g., 6+6, 6+5).

**Figure 2.**
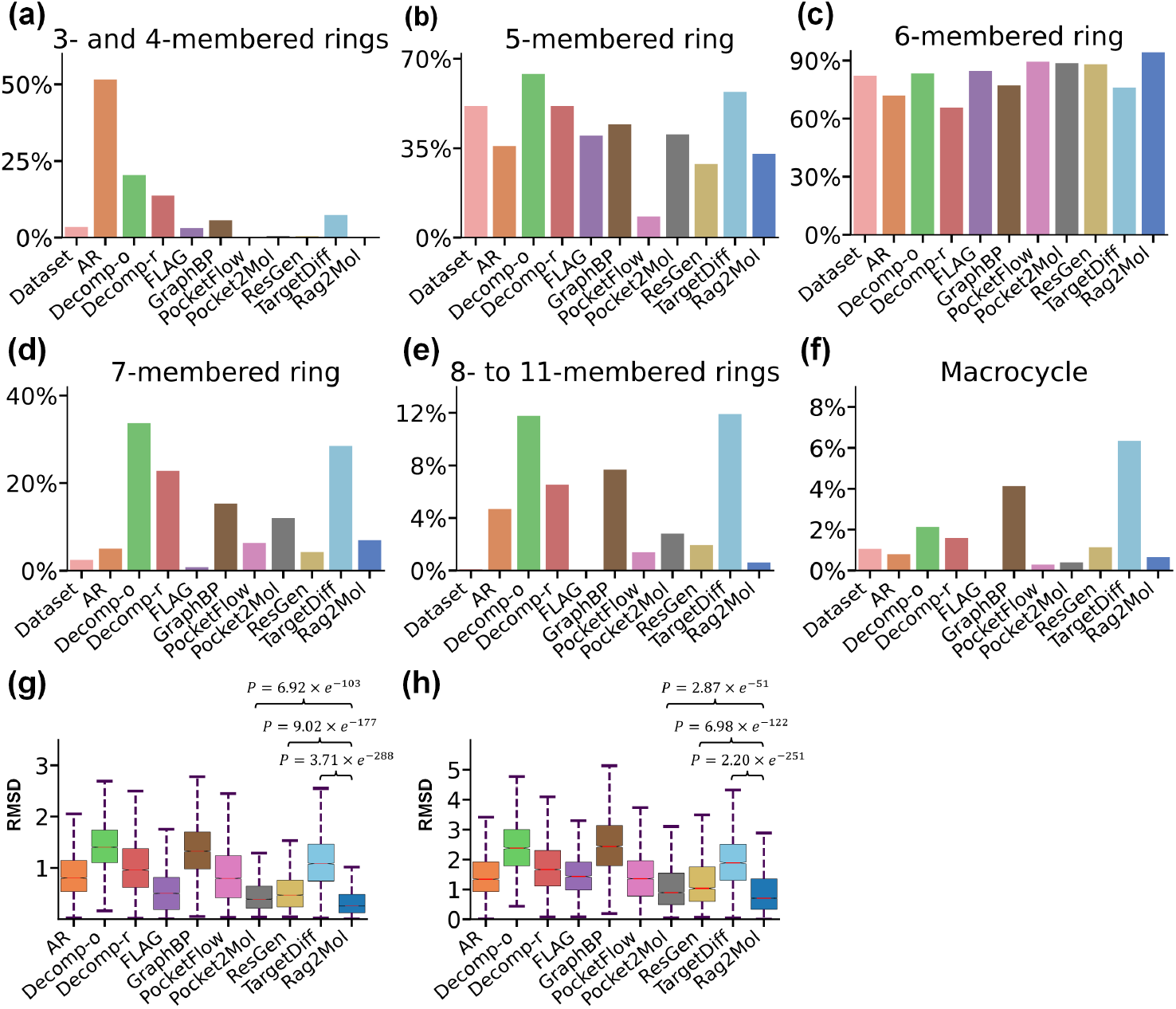
Conformational quality assessment. (a-f) **Rings analysis.** We compare the ratio of rings of different sizes in generated molecules by different SBDD models. The statistics of native molecules in the dataset are shown in the first column. (g-h) **SBDD models’ conformational capability**. The RMSD offsets between the generated molecular structures and the structures calculated by RDKit^53^ and QVina^54^ are shown, respectively.

Figure 2g shows that Rag2Mol generates smoother conformations, reflected by the lowest RMSD values, while Figure 2h indicates Rag2Mol captures complex geometric distributions within pockets more effectively than other methods. Statistically significant p-values further support these findings.

We also evaluated 9 common bond angles and dihedral angles using KL divergence, where lower values signify higher consistency with the test set. As shown in Figure 3, Rag2Mol achieved the best performance in 5 metrics and competitive results in the remaining ones, particularly excelling in dihedral angles, a challenging metric for other SBDD methods. We believe that the introduction of retrieval augmentation has mitigated SBDD’s limitations in modeling complex substructures.

**Figure 3.**
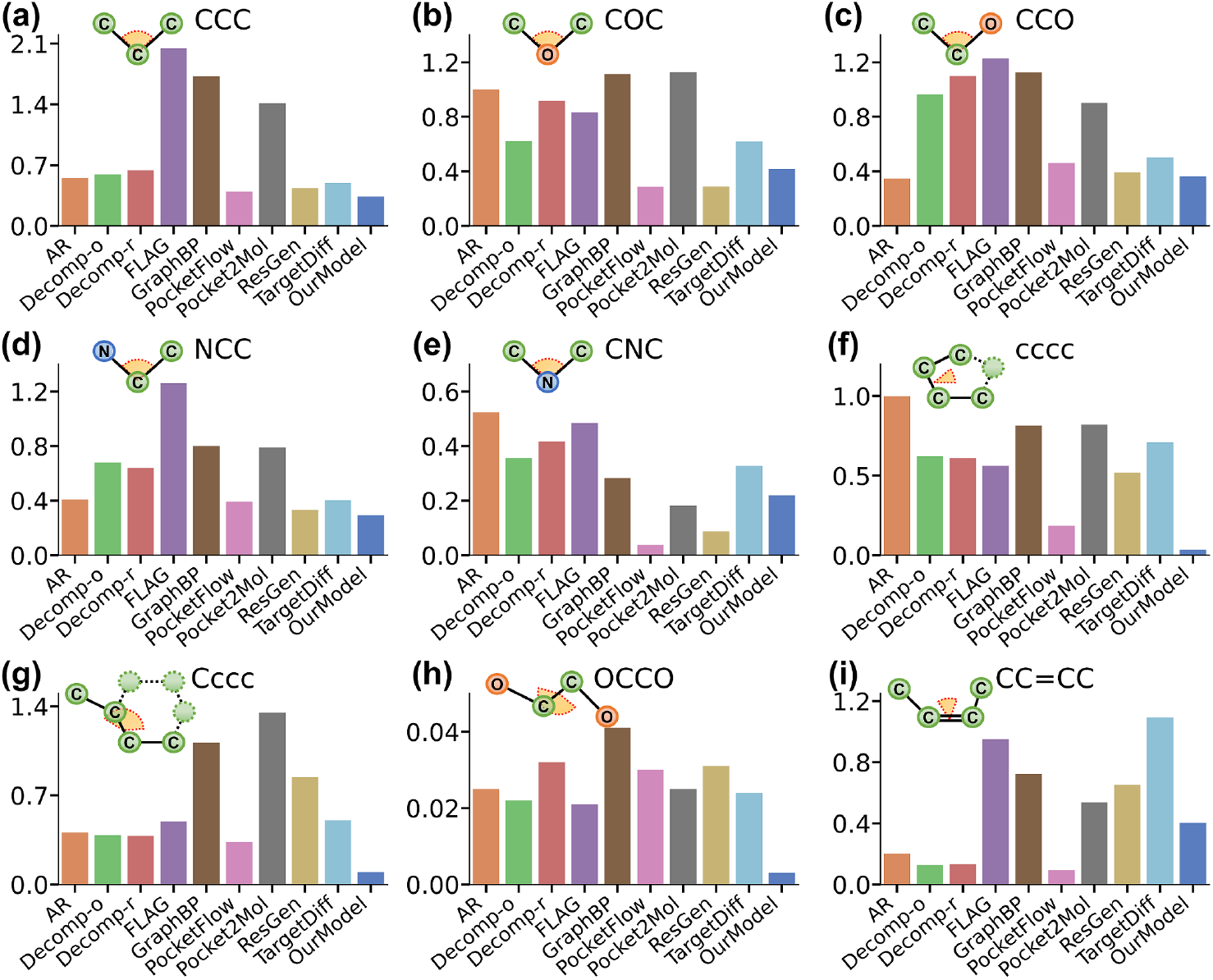
Bond angles and dihedral angles analysis. The distributions of the bond angles and dihedral angles of the generated molecules agree with the test set by using the KL divergence. We provide a simple diagram for each type of angles.

### 3.3 Interaction pattern analysis

Figure 4b demonstrates Rag2Mol’s ability to capture microscopic interaction patterns on therapeutic target: AKT1 (PDB id: 4gv1), due to its well-studied binding patterns. Rag2Mol-generated molecules display reasonable binding poses, suggesting effective inference of hit positioning within the protein pocket. Using PLIP^55^, we analyze interactions between the target and Rag2Mol-generated ligand, comparing them to experimentally validated active ligand. Rag2Mol reproduces most key interactions observed in experimental ligands (6/7 for 4gv1), including hydrophobic interactions with ALA177 and LYS179 and hydrogen bonds with ALA230, GLU234, and ASP292. Importantly, Rag2Mol introduces 4 extra, physically plausible interactions that further enhance binding potential. Moreover, we analyze two more targets (CDK2 and AROK); the detailed interaction statistics and visualizations are provided in sections 6 and 7 of Supplementary Information.

**Figure 4.**
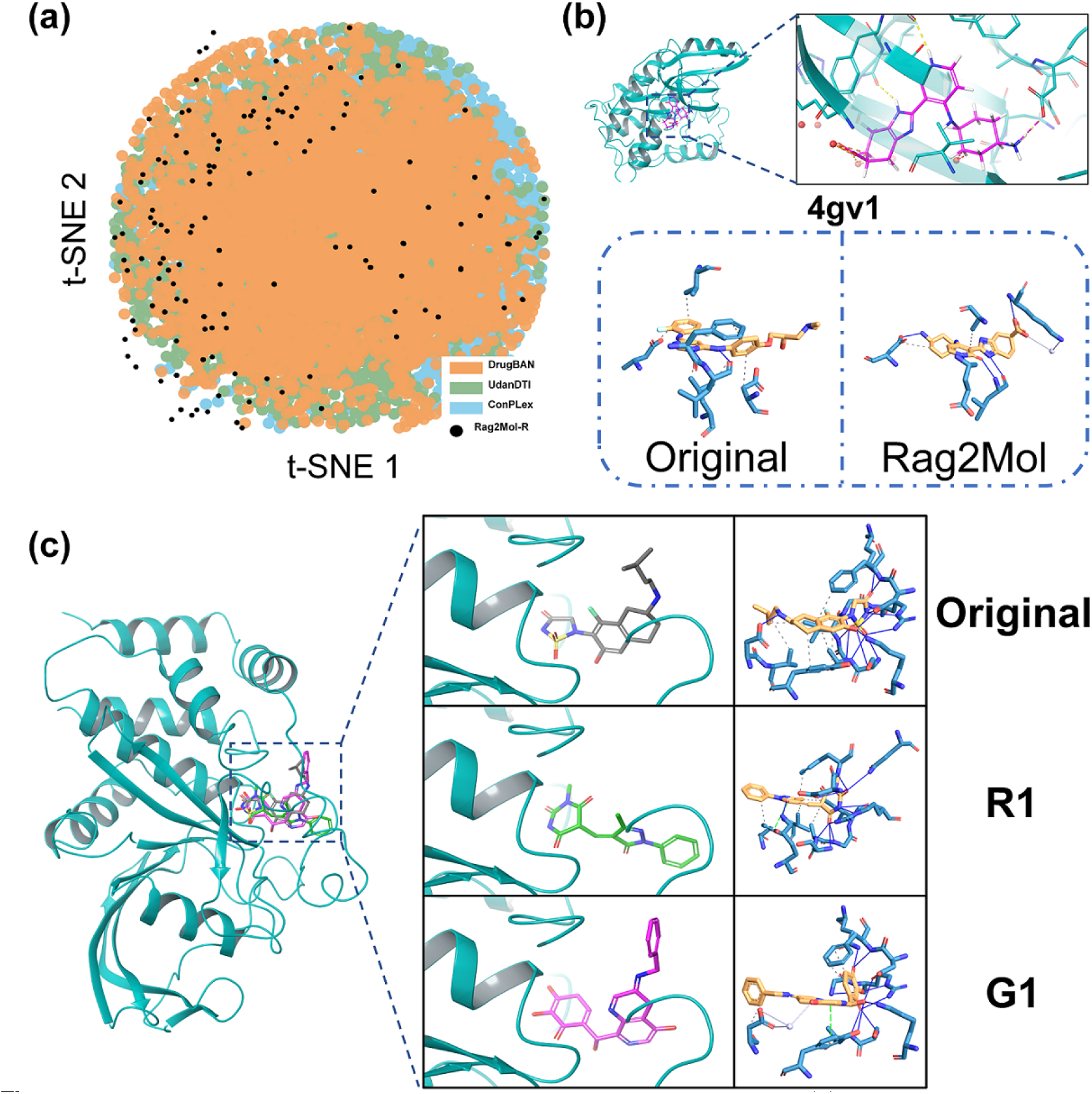
The application results of Rag2Mol on different real-world cases. (a) **Chemical space map representations for compounds screened by different models**. t-SNE is used for visualizing data by giving each data point a location in this two-dimensional map. The screening hits from Rag2Mol are represented in black. In comparison, compounds screened by DrugBAN, UdanDTI, and ConPLex are depicted in orange, green, and blue, respectively. (b) **Protein–ligand interaction analysis**. The top quadrants show the poses of Rag2Mol’s generated ligands within the protein pocket, whereas the below quadrants denote the protein-ligand interaction patterns for the original and generated ligands, respectively. **(C) Rag2Mol on PTPN2**. The left image shows the overlap of the three drugs within the protein pocket, while the two columns on the right display their binding conformations and binding interaction analysis.

### 3.4 Detecting similar lead compounds

In Table 2, we compared the molecules identified by Rag2Mol-R with those by advanced virtual screening models. Overall, Rag2Mol-R demonstrated superior performance across nearly all metrics related to binding affinity and drug-likeness. It not only inherits the high affinity of designed molecules of Rag2Mol (we demonstrate the connection in Supplementary Figure S2) but also ensures the identified drug candidates are readily synthesizable. Note that a significant percentage of molecules identified by Rag2Mol-R (approximately 74%) showed higher affinity scores than the native ligands on the target, obviously outperforming other virtual screening models.

**Table 2.** The mean binding energies and drug-likeness properties of top 1/3/5/10 molecules in drug repurposing. (*↑*)/ (*↓*) indicates larger / smaller is better. Top 1 results are highlighted with **bold**.

|  | Test set | UdanDTI | DrugBAN | ConPLex | Our model |
| --- | --- | --- | --- | --- | --- |
| Affinity<sup>1</sup>(↑) | - | 52.8% | 53.0% | 57.4% | **74.2%** |
| **Top 1 / Top 3** |  |  |  |  |  |
| Vina Dock(↓) | -7.204 / -7.204 | -9.001 / -8.384 | -8.100 / -7.567 | -10.141 / -9.837 | **-10.625 / -10.411** |
| QED(↑) | 0.476 / 0.476 | 0.351 / 0.355 | 0.272 / 0.282 | 0.365 / 0.393 | **0.509 / 0.503** |
| SA(↑) | 0.728 / 0.728 | 0.610 / 0.604 | 0.591 / 0.594 | 0.714 / 0.727 | **0.814 / 0.823** |
| Lipinski(↑) | 4.340 / 4.340 | 3.990 / 4.020 | 3.910 / 3.853 | 4.240 / 4.300 | **4.990 / 4.963** |
| LogP | 0.894 / 0.894 | 4.520 / 4.618 | 2.269 / 4.026 | 5.179 / 4.766 | 4.463 / 4.313 |
| **Top 5 / Top 10** |  |  |  |  |  |
| Vina Dock(↓) | -7.204 / -7.204 | -8.312 / -8.215 | -7.252 / -6.326 | -9.635 / -9.281 | **-10.273 / -10.047** |
| QED(↑) | 0.476 / 0.476 | 0.352 / 0.361 | 0.290 / 0.306 | 0.407 / 0.438 | **0.500 / 0.507** |
| SA(↑) | 0.728 / 0.728 | 0.622 / 0.638 | 0.598 / 0.597 | 0.733 / 0.741 | **0.822 / 0.825** |
| Lipinski(↑) | 4.340 / 4.340 | 4.040 / 4.124 | 3.830 / 3.823 | 4.336 / 4.379 | **4.962 / 4.962** |
| LogP | 0.894 / 0.894 | 4.585 / 4.510 | 3.761 / 3.695 | 4.643 / 4.394 | 4.174 / 4.060 |

Furthermore, we used three baseline models to search approximately 10,000 molecules for a randomly selected target protein (PDB: 3zkg) and visualized the chemical space of those molecules using the t-SNE algorithm. As illustrated in Figure 4a, even when existing advanced virtual screening models are prompted to yield a high number of drug candidates, they fail to encompass the top 115 valuable compounds identified by Rag2Mol-R. This effectively highlights a broader coverage of the chemical landscape and the precise targeting capability of Rag2Mol-R. Finally, the retrieval database used by Rag2Mol and the similarity search database utilized by Rag2Mol-R are both scalable.

### 3.5 Case study: PTPN2

Protein tyrosine phosphatases PTPN2 and PTPN1 are central regulators of inflammation, and their deletion in tumor or immune cells enhances anti-tumor immunity. However, phosphatases have long been considered undruggable, for they have highly polar active sites. Recently, AC484^56^ was identified as a promising monotherapy.

In response to this challenge, we applied Rag2Mol’s two workflows to this complex problem. In Glide’s precise docking evaluation, the drug candidates G1 and R1 identified by Rag2Mol-G and Rag2Mol-R exhibit affinity scores of -13.8 and -12.1, respectively, surpassing AC484’s score of -11.2. Additionally, G1 and R1 both demonstrate higher synthetic accessibility, while maintaining comparable molecular weights and logP values. As illustrated in Figure 4c, G1 shares structural similarities with AC484, while R1 presents a distinct molecular scaffold. All three molecules exhibit significant overlap within the target pocket.

Interaction analysis shows that R1 and G1 capture most of the key hydrogen bonds formed with LYS122, ASP182-PHE183, SER217-ARG222, and GLN264, which play a crucial role in AC484’s inhibition of phosphatases. The hydrophobic interactions of R1 and G1 closely match those of AC484. Additionally, both R1 and G1 form extra pi-stacking interactions with the TYR48’s side chain, further strengthening their binding. Notably, G1 enhances stability by converting the original hydrophobic interaction between TYR48 and AC484 into a hydrogen bond. Complete statistical metrics are provided in the Supplementary Table S8.

## 4 Conclusion

This paper introduces a drug discovery protocol with two distinct workflows. Inspired by the concept of RAG, we developed Rag2Mol, a RAG-based E(3)-equivariant GNN generative model. Rag2Mol actively integrates retrieved reference knowledge during both training and generation procedures to generate 3D drug-like molecules targeting protein pockets. Experimental results demonstrate that introducing retrieved knowledge enables the SBDD model to better comprehend biochemical rules and perceive the geometric environment, thereby meeting the demands of real-world drug design projects. The creation of Rag2Mol-G and Rag2Mol-R reflects a strategic decision to address different drug discovery scenarios. Rag2Mol-G focuses on generating novel molecules with optimal properties, while Rag2Mol-R prioritizes the identification of readily available candidates, both serving a unique role in the drug discovery pipeline. The components of our protocol, including the two-level retriever, docking module, databases, similarity search module, and filtering mechanisms, all exhibit strong scalability. We believe this work offers valuable new perspectives for the drug discovery field.

