## Supplementarty Information for "Rag2Mol: Structure-based drug design based on Retrieval Augmented Generation"

### Supplementary Information

### Rag2Mol: Retrieval augmentation assists structure-based molecular generation in drug discovery

### 1 Dataset, databases and evaluation

#### 1.1 CrossDock dataset

Rag2Mol and other structure-based drug design (SBDD) baselines are trained and tested on CrossDock [1] dataset, which is designed to enhance SBDD models by providing a large collection of protein-ligand complexes. This dataset specifically includes cross-docked protein-ligand pairs, which are generated by docking ligands to multiple protein targets. The primary aim is to facilitate the training and evaluation of 3D convolutional neural networks (CNNs) for predicting binding affinities and improving docking accuracy. The dataset offers a diverse set of conformations, enhancing the generalizability of models developed for drug discovery applications. Following [2, 3], the protein sequence identity between the test set and training set is constrained to be less than 30%, resulting in about 100,000 training pairs and 100 test proteins. For each pocket sample on the test set, each SBDD model generates about 100 molecules. Notably, we compare the screening task on the same test set. We provide the gene ids of test set in Table S1.

Table S1. Gene ids for test set.

| Name list | Total |
|---|---|
| PHP_SULSO, CHIB_SERMA, MENE_BACSU, DPP2_HUMAN, NQO1_HUMAN, CHIB_SERMA, UPPS_ECOLI, SIR3_HUMAN, RIBB_VIBCH, AROE_THET8, RG1_RAUSE, PLCD1_RAT, TNKS1_HUMAN, BGL07_ORYSJ, CPXB_BACMB, GUX1_HYPJE, SQHC_ALIAD, BTRN_BACCI, BACE2_HUMAN, CHOD_BREST, F16P1_HUMAN, NPD_THEMA, POL_FOAMV, ODBB_THET8, PTGIS_HUMAN, PTGIS_HUMAN, LMBL1_HUMAN, ACE_HUMAN, BAPA_SPHXN, NEP_HUMAN, CCPR_YEAST, M3K14_HUMAN, MCCF_ECOLX, PAK4_HUMAN, CPXB_BACMB, BSD_ASPTE, ABL2_HUMAN, DYRK2_HUMAN, ACE_HUMAN, NOS3_HUMAN, BACE2_HUMAN, AROE_THET8, PYRD_TRYCC, ATS5_HUMAN, SDIA_ECOLI, FKB1A_HUMAN, CDK6_HUMAN, PAK4_HUMAN, PA21B_PIG, IMA1_HUMAN, AK1BA_HUMAN, AKT1_HUMAN, NOS1_HUMAN, CDK6_HUMAN, P2Y12_HUMAN, CHIB_SERMA, NOS1_HUMAN, KS6A3_HUMAN, FKB1A_HUMAN, NR1H4_HUMAN, NAGZ_VIBCH, ATS5_HUMAN, DFPA_LOLVU, DIDH_RAT, PLCD1_RAT, NQO1_HUMAN, OLIAC_CANSA, HDAC8_HUMAN, OLIAC_CANSA, NOS2_HUMAN, DYRK2_HUMAN, CDK6_HUMAN, IPMK_HUMAN, F16P1_HUMAN, PHP_SULSO, NAGZ_VIBCH, NEP_HUMAN, NOS3_HUMAN, EFTU1_ECOLI, NOS2_HUMAN, DHAK_ECOLI, CHIA_SERMA, BSD_ASPTE, TRAR_RHIRD, PPIA_HUMAN, TBK1_HUMAN, IDHP_HUMAN, PAC_ECOLX, BAZ2A_HUMAN, NOS3_HUMAN, ODBB_THET8, CHIB1_ASPFM, VAOX_PENSI, IPMK_HUMAN, PYRD_TRYCC, NOS3_HUMAN, BGL07_ORYSJ, NQO1_HUMAN, TIAM1_HUMAN, HMD_METJA, BTRN_BACCI, PNTM_STRAE, DHAK_ECOLI, CONA_CANCT, NR1H4_HUMAN, M3K14_HUMAN, PA2B8_DABRR, GSTP1_HUMAN, CD38_HUMAN, TIAM1_HUMAN, LAT_MYCTU, GUX1_HYPJE, GSTP1_HUMAN, UBE2T_HUMAN, TNKS2_HUMAN, DIDH_RAT, ABL2_HUMAN, LMBL1_HUMAN, BGAT_HUMAN, COTA_BACSU, TRAR_RHIRD, DPO4_SULSO, TNKS1_HUMAN, PAC_ECOLX, TNKS2_HUMAN, HDHA_ECOLI, RG1_RAUSE, DFPA_LOLVU, P2Y12_HUMAN, IDHP_HUMAN, CCPR_YEAST, DPP2_HUMAN, NQO1_HUMAN, CHOD_BREST, NOS1_HUMAN, SQHC_ALIAD, IMA1_HUMAN, UBE2T_HUMAN, PA2B8_DABRR, AKT1_HUMAN, COAA_MYCTU, CHIA_SERMA, POL_FOAMV, TBK1_HUMAN, CAT_ECOLX, EFTU1_ECOLI, GLMU_STRPN, ROCO4_DICDI, PNTM_STRAE, HDHA_ECOLI, GLMU_STRPN, LMBL1_HUMAN, CDK6_HUMAN, AK1BA_HUMAN, CAT_ECOLX, LAT_MYCTU, QPCT_HUMAN, CD38_HUMAN, KS6A3_HUMAN, XANLY_BACGL, NOS1_HUMAN, SIR3_HUMAN, QPCT_HUMAN, PYRE_BACAN, EXG1_CANAL, BGAT_HUMAN, UPPS_ECOLI, Y635_MYCTU, PPIA_HUMAN, CHIB_SERMA, CHIB1_ASPFM, MURA_ECOLI, XANLY_BACGL, MCCF_ECOLX, Y635_MYCTU, PA21B_PIG, VAOX_PENSI, GRK4_HUMAN, PYRD_TRYCC, PYRE_BACAN, SDIA_ECOLI, HMD_METJA, EXG1_CANAL, HDAC8_HUMAN, CONA_CANCT, GRK4_HUMAN, ROCO4_DICDI, BAPA_SPHXN, PHKG1_RABIT, PYRD_TRYCC, NPD_THEMA, BAZ2A_HUMAN, PHKG1_RABIT, MENE_BACSU, COTA_BACSU, MURA_ECOLI, DPO4_SULSO, RIBB_VIBCH, COAA_MYCTU, LMBL1_HUMAN | 93 |

#### 1.2 Databases

We build two molecular compound databases for different goals. During the retrieval step (Figure

1a), we need to apply rapid virtual screening on small molecules that are more likely to interact with the target proteins. Thus we choose the combination of BindingDB and PDBBind.

For the similarity search step, the database used must cover a broad chemical landscape, ensuring diversity in chemical scaffolds, and guaranteeing synthesizability. Based on these criteria, we created a database by selecting purchasable drug-like molecules from the most extensive existing bioactive molecular databases, including GEOM, ChEMBL, and ZINC15, followed by a deduplication process. Here is the detailed explanation:

1. BindingDB [4] is a public database that contains experimentally determined binding affinities for protein-ligand interactions. It provides a comprehensive resource for researchers to access quantitative data on small molecule interactions with various biological targets, aiding in the development of new drugs. We use all purchasable compounds of having affinity better than 10 uM in this database.
2. PDBBind [5] is a dataset that integrates binding affinity data with structural information from the Protein Data Bank (PDB). It features a curated collection of protein-ligand complexes, providing a valuable resource for evaluating and developing computational models for predicting binding affinities in structure-based drug design. We collect the ligands from the complexes in version 2020 of PDBBind.
3. GEOM [6] is a database with 37 million molecular conformations annotated by energy and statistical weight for over 450,000 molecules. It is a database focused on geometric properties of protein-ligand interactions. It offers a collection of molecular geometries and conformational data, facilitating the analysis of molecular shapes and binding modes, which are critical for understanding interactions in drug discovery. We use the drug-like molecules from the initial version (data relating to SARS-CoV-2 are not included).
4. ChEMBL [7] is a manually curated database of bioactive molecules with drug-like properties. It brings together chemical, bioactivity and genomic data to aid the translation of genomic information into effective new drugs. It serves as a vital resource for medicinal chemists and researchers in drug discovery, providing data for compound efficacy, safety, and pharmacological profiles. We choose all distinct compounds in this database.
5. ZINC15 [8] is a free database of commercially available compounds for virtual screening. It contains a vast collection of drug-like molecules, providing researchers with a resource for identifying potential drug candidates. The database is designed to facilitate the selection of compounds for experimental testing and structure-based drug design. We choose compounds that are available for purchase with LogP between -2 and 6 and molecular weight between 150 and 550.

Table S2. Database statistics.

| Database | Drugs |
|---|---|
| BindingDB | 30, 827 |
| PDBBind | 13, 029 |
| GEOM | 437, 572 |
| ChEMBL | 2, 372, 524 |
| ZINC15 | 4, 014, 216 |

#### 1.3 Evaluation

The three design criteria for a well-designed SBDD model are: (1) Fundamentally, the molecules generated by the model should be capable of tightly binding to the target protein pocket. Thus SBDD model requires to learn the biochemical information within the given pocket while considering the topological distribution and interaction patterns, ultimately generating drug candidates with greater binding potential. (2) SBDD model could predict reasonable atom positions and chemical bonds based on the surrounding topological environment. Molecules generated by SBDD model should exhibit physically plausible conformations and valid geometric distributions within pocket. (3) The molecules should also exhibit favorable intrinsic properties, such as drug-likeness, synthetic accessibility, and structural diversity, which reflects the SBDD model's generalization.

To comprehensively assess and compare vinilla Rag2Mol against existing SOTA SBDD models with respect to these three design criteria, we introduced the following evaluation scheme. To assess the first and third design criterias, we compared molecules generated by Rag2Mol and eight other baselines on the publicly available CrossDock dataset, using widely recognized metrics: (a) Vina Dock and Vina Score estimate the binding affinity between generated molecules and protein pockets before and after docking, respectively. (b) $Affinity^1$ and $Affinity^2$ calculate the percentage of molecules with higher affinities to the existing ligands before and after docking, respectively. (c) QED is a drug-likeness metric, quantitatively estimating how likely a molecule is a potential drug. (d) SA is the synthetic accessibility score, with higher values indicating easier synthesis (standardized between 0 and 1). (e) Lipinski counts the number of Lipinski's rules obeyed by current molecule, reflecting empirical laws of drug-likeness. (f) LogP represents the octanol-water partition coefficient, with a range between -0.4 and 5.6 for an ideal drug. (g) Diversity is calculated as the average Tanimoto similarities of the generated molecules for the individual pockets. The relevant binding metrics are derived using QVina [9] and the chemical properties are calculated by RDKit (https://www.rdkit.org/).

To evaluate the second design criteria, we compared the distributions of bond angles and dihedral angles of the generated molecules agree with the test set using Kullback-Leibler (KL) divergence. The ratio of ring-structures with different sizes is also analyzed. Moreover, we also quantitatively compared conformation quality by extracting 1D sequence information (SMILES) from the generated molecules and re-predicting their conformations using both energy-based chemical software (RDKit) and deep learning-based docking software (FABind [10]), separately. The root-mean-square deviation (RMSD) was used to measure pose differences before and after re-prediction. For interaction pattern analysis, we randomly selected several native cases for Rag2Mol and extracted their protein-ligand interaction graphs. Finally, for selected molecules from real compound libraries, the same metrics are used to evaluate Rag2Mol-R and other SOTA virtual screen methods.

#### 2 Baselines

**SBDD baselines:**

1. **GraphBP** [11] employs graph neural networks to represent protein pockets and ligands as graphs, capturing complex atomic relationships and interactions. This graph-based approach allows the model to learn effective structural representations, resulting in enhanced accuracy and efficiency in ligand generation.
2. **Pocket2Mol** [12] generates ligands by leveraging the three-dimensional geometry of protein binding pockets. It samples molecular conformations that fit within the spatial constraints of the pocket, ensuring compatibility and high binding affinity. The model uses a probabilistic framework for diverse candidate generation directly from pocket structures.
3. **ResGen** [13] adopts a multi-scale modeling approach to create ligands by considering both the binding pocket and the surrounding residues. This model captures interactions at varying resolutions, which enhances the understanding of ligand-receptor dynamics and allows for tailored ligand design that improves binding affinity and specificity.
4. **AR** [2] model generates molecules sequentially, predicting each atom based on previously generated atoms and the context of the binding site. This autoregressive process incorporates both local and global structural information, facilitating the creation of chemically viable conformations tailored to the target pocket.
5. **FLAG** [14] uses a fragment-based strategy to generate ligands by assembling small chemical fragments. It captures interactions within the protein pocket while leveraging existing fragment libraries, allowing for efficient chemical space exploration and maintaining high binding potential.
6. **TargetDiff** [15] focuses on identifying potential binding sites by analyzing structural differences among related target proteins. This model generates ligands targeting conserved binding regions, enhancing the discovery of new hits for difficult targets by leveraging evolutionary conservation.
7. **Decompdiff** [16] breaks down complex molecular structures into simpler components, focusing on individual ligand parts and their interactions with the pocket. By recombining these components, it generates diverse ligands while ensuring compatibility with the binding site.

**Virtual screen baselines:**

1. **ConPLex** [17] leverages contrastive learning in a protein language space to predict drug-target interactions by encoding protein sequences and ligand features into a shared embedding space. It trains the model to distinguish between positive (interacting) and negative (non-interacting) pairs, enhancing the representation of meaningful interactions. By utilizing both protein and ligand embeddings, ConPLex effectively captures complex relational patterns, leading to improved predictions of binding affinities and interaction probabilities. This approach allows for a nuanced understanding of the interaction landscape between drugs and their protein targets.
2. **DrugBAN** [18] utilizes a bidirectional attention mechanism to improve the prediction of drug-target interactions. By integrating drug and target embeddings, it focuses on relevant features and interactions from both entities, allowing for a more nuanced understanding of their relationship. The attention layers help highlight critical features that contribute to binding affinity and specificity.

3. **UdanDTI** [19] is designed to predict drug-target interactions using an unbalanced dual-branch framework that simultaneously considers multiple biological contexts. This model leverages large language model embeddings and attentive module to enhance its predictive capabilities. By explicitly fusing the encoded information from protein and drug, UdanDTI improves the robustness of interaction predictions.

### 3 Formulas for predictors and loss functions

#### 3.1 Predictors

We use $(v_f, \boldsymbol{v}_f)$ to represent the extracted hidden representations, which capture not only the chemical and geometric attributes within protein pocket, but also the generally molecular structural laws and complete drug-target interaction patterns. These representations are used for focal atom predictor, position predictor and element-and-bond predictor.

**Focal atom predictor**

As shown in Figure 1b, the first subtask is to predict the focal atom as the starting seed among generated atoms (protein pocket atoms when initialization). Specifically, the probability of $i$th atom to be a focal atom $p_f{}^{(i)}$ could be calculated as follow:

$$\left(p_f{}^{(i)}, \boldsymbol{p}_f{}^{(i)}\right) = G_f\left(v_f{}^{(i)}, \boldsymbol{v}_f{}^{(i)}\right), \tag{1}$$

$$\hat{p}_f{}^{(i)} = \sigma_{sg}\left(p_f{}^{(i)}\right), \tag{2}$$

where $\sigma_{sg}$ is the sigmoid activation function.

**Position predictor**

Researchers have employed mixture density network to model the spatial distribution of atoms, effectively avoiding the potential accuracy risks associated with directly generating atomic coordinates. Additionally, transforming the regression task to a generative one enhances both the robustness and diversity of the model. Therefore, we also assume that the interatomic distance distributions conform to a multivariate Gaussian mixture distribution with diagonal covariance, that is, $p(\Delta r) = \sum_{k=1}^{K} \pi_k \cdot N(\mu_k, \Sigma_k)$. Where $K$ indicates the number of Gaussians and $\Delta r$ represents the relative coordinates toward the focal atom. $\pi$, $\mu$, $\Sigma$ are the combination coefficients, mean and variance of the multivariate Gaussian mixture distribution, respectively. Assume that the focal atom is the $i$th atom, the distribution of newly generated atom position could be calculated as follow:

$$\left(\mu^{(i)}, \boldsymbol{\mu}^{(i)}\right) = G_\mu\left(v_f{}^{(i)}, \boldsymbol{v}_f{}^{(i)}\right), \tag{3}$$

$$\left(\Sigma^{(i)}, \boldsymbol{\Sigma}^{(i)}\right) = G_\Sigma\left(v_f{}^{(i)}, \boldsymbol{v}_f{}^{(i)}\right), \tag{4}$$

$$\left(\pi^{(i)}, \boldsymbol{\pi}^{(i)}\right) = G_\pi\left(v_f{}^{(i)}, \boldsymbol{v}_f{}^{(i)}\right), \tag{5}$$

$$\left(\hat{\mu}^{(i)}, \hat{\Sigma}^{(i)}, \hat{\pi}^{(i)}\right) = \left(\boldsymbol{\mu}^{(i)}, exp\left(\boldsymbol{\Sigma}^{(i)}\right), \sigma_{sf}\left(\pi^{(i)}\right)\right). \tag{6}$$

Where $\sigma_{sf}$ is the softmax function. We could sample the next atom in 3D space from the predicted multivariate Gaussian mixture distribution based on the focal atomic coordinates.

**Element-and-bond predictor**

Once we have sampled the position of next atom, the element-and-bond predictor will predict the element type and the valence bonds with existing atoms in molecular fragment of newly generated atom. Similar to Message passing module, assume that the $i$th atom is newly generated, the information of k-nearest atoms would be aggregated on the sampled position:

$$\left(z^{(i)}, \boldsymbol{z}^{(i)}\right) = \sum\nolimits_{KNN} G_a\left(v_f{}^{(i)}, \boldsymbol{v}_f{}^{(i)}, e_{f\rightarrow i}, \boldsymbol{e}_{f\rightarrow i}\right). \tag{7}$$

Thus we could calculate the distribution of pre-set element types:

$$\left(p_e{}^{(i)}, \boldsymbol{p}_e{}^{(i)}\right) = G_e\left(z^{(i)}, \boldsymbol{z}^{(i)}\right), \tag{8}$$

$$\hat{p}_e{}^{(i)} = p_e{}^{(i)}. \tag{9}$$

Besides, unlike other SBDD works predicting possible bonding by chemical rules, a specific predictor is trained to predict the bonding relationship between the $i$th atom and existing atoms. Take the $j$th atom as an example, the edge features between atom $i$ and $j$ could be embedded as $\left(e^{(i,j)}, \boldsymbol{e}^{(i,j)}\right)$ following formula (18). And we represent and encoded the potential bonding relationship as follows:

$$\left(z^{(i,j)}, \boldsymbol{z}^{(i,j)}\right) = G_{b1}\left(\left[z^{(i)}; v^{(j)}; e^{(i,j)}\right], \left[\boldsymbol{z}^{(i)}; \boldsymbol{v}^{(j)}; \boldsymbol{e}^{(i,j)}\right]\right). \quad (10)$$

Where $[;]$ is the concatenating operation. Then the probability of bonding type between atom $i$ and $j$ is calculated by a trigonometry self-attention, which is proposed by [12] and has been proved efficiency in capturing chemical bonding relationship.

$$\left({p_b}^{(i,j)}, {\boldsymbol{p}_b}^{(i,j)}\right) = G_{b2}\left(Attn_t\left(z^{(i,j)}, \boldsymbol{z}^{(i,j)}\right)\right), \quad (11)$$

$$\hat{p}_b{}^{(i,j)} = {p_b}^{(i,j)}. \quad (12)$$

#### 3.2 Loss functions

Here we provide the detailed formulas of loss functions:

$$\mathcal{L}_{focal} = -\frac{1}{n}\sum_{i=1}^{n} \hat{p}_f{}^{(i)} \times log\left({p_f}^{(i)}\right) + \left(1 - \hat{p}_f{}^{(i)}\right) \times log\left(1 - {p_f}^{(i)}\right) \quad (24)$$

$$\mathcal{L}_{pos} = \frac{1}{n}\sum_{i=1}^{n} -log \sum_{k=1}^{K} \hat{\pi}_k^{(i)} \cdot N\left(\hat{\mu}_k^{(i)}, \hat{\Sigma}_k^{(i)}\right) \quad (25)$$

$$\mathcal{L}_{element} = -\frac{1}{n}\sum_{i=1}^{n} \hat{p}_e{}^{(i)} \times log\left({p_e}^{(i)}\right) \quad (26)$$

$$\mathcal{L}_{bond} = -\frac{1}{n}\sum_{i=1}^{n}\sum_{j \in fragment} \hat{p}_b{}^{(i,j)} \times log\left({p_b}^{(i,j)}\right) \quad (27)$$

### 4 Implementation, tools, and other set-ups

#### 4.1 Implementation of workflows

We have developed two workflows for drug discovery: Rag2Mol-G and Rag2Mol-R. In the Rag2Mol-G workflow, following the widely accepted threshold settings, we filter the generated molecules based on the following criteria: $\mathbf{Vina} \in [-20,-5], \mathbf{QED} \in [0.5,2], \mathbf{SA} \in [0.5,2], \mathbf{Lipinski} \in [4,5],$ and $\mathbf{LogP} \in [0,4]$. The filtered drug candidates are then subjected to precise binding affinity calculations and subsequent wet-lab experiments.

In the Rag2Mol-R workflow, after applying the same filtering criteria, sphere exclusion clustering is applied on the molecules by RDKit. Following the recommendations in [20], a minimum distance threshold of 0.6 between cluster centroids is employed, with Morgan fingerprints serving as the clustering feature. From each cluster, representative molecule is randomly selected as scaffold template. Based on these templates, we search for the top 4 similar molecules within existing synthesizable compounds. These molecules are deduplicated and then subjected to accurate docking software Glide [21], yielding the final set of drug candidates.

#### 4.2 Tools

We use several tools in this study:

1. **QVina** is an open-source docking tool designed for high-throughput virtual screening, with speeding up AutoDock Vina software. Qvina is known for its efficiency in handling large compound libraries while maintaining a balance between speed and prediction quality. All the metrics related to docking are calculated by QVina.
2. **Glide** is a precise molecular docking software that employs a multi-step approach to predict the binding of small molecules to protein targets. It utilizes a scoring function based on empirical data, which balances accuracy and computational efficiency. Glide also incorporates flexible ligand and receptor docking, enabling it to explore conformational changes that may occur during binding. In this study, it is used for high-precision docking in real cases.
3. **FABind** is a deep learning-based docking method that emphasizes speed and accuracy in predicting protein-ligand binding interactions. It leverages a fragment-based approach, which allows for efficient exploration of the binding pocket by generating ligand conformations from smaller molecular fragments. FABind's advanced scoring function integrates both empirical data and detailed interaction models, resulting in improved binding affinity predictions while significantly reducing computational time. We use it to construct extra molecular knowledge for its speed.
4. **RDKit** is an open-source cheminformatics toolkit designed for cheminformatics and machine learning applications. It provides a comprehensive suite of functionalities, including molecular manipulation, descriptor calculation, and substructure searching. RDKit supports various file formats for input and output, enabling seamless integration with other data analysis tools. Additionally, it is widely used in the development of predictive models for drug discovery and molecular design.

#### 4.3 Other set-ups

**Molecular similarity**. All the molecular similarity is calculated by the Tanimoto coefficient using RDKit, which is defined as follows:

$$\mathrm{T}(A,B) = \frac{|A \cap B|}{|A \cup B|} = \frac{|A \cap B|}{|A| + |B| - |A \cap B|}, \quad ( )$$

Where $A$ and $B$ are the Morgan fingerprints of different molecules.

**RMSD** quantitatively compares the differences within two molecular structures, which is calculated as:

$$\mathrm{RMSD}(M_1, M_2) = min_\theta \sqrt{\frac{1}{N}\sum_{i=1}^{N} \left\| M_1^{(\mathrm{i})} - M_2^{(\mathrm{i})} \right\|^2}, \quad ( )$$

Where $M_1$ and $M_2$ are the coordinates of different molecules, $N$ is the number of heavy atoms and $\theta$ indicates the alignment function.

**KL divergence** can be used to measure the degree of difference between two distributions. Assume that $\mathrm{p}(x)$ and $\mathrm{q}(x)$ are two probability distributions for the discrete variable $x$, $\mathrm{KL}(p \parallel q)$ is defined as follows:

$$\mathrm{KL}(p \parallel q) = \sum_{i=1}^{n} p(x_i) \cdot log \frac{p(x_i)}{q(x_i)}. \quad ( )$$

Besides, we provide the hyperparameters for implementation.

Table S3. Hyperparameter setting.

| Module | Parameters | values |
|---|---|---|
| Feature extractor | Scalar hidden channels | 256 |
| | Vector hidden channels | 64 |
| Encoder | Layers | 6 |
| | Knn | 48 |
| Message-passing | Scalar attn heads | 16 |
| | Vector attn heads | 8 |
| | Cross-Knn | 48 |
| Training | Learning rate | 0.0002 |
| | Batch size | 6 |
| | Optimizer | Adam |
| | Epochs | 200 |
| Sampling | Pocket-specific database size | 5000 |
| | Vina threshold | 0.0 |
| | Random choose nums | 64 |
| | Beam size | 300 |
| | Max steps | 50 |

### 5 Complete comparison between Rag2Mol and baselines

Table S4. Complete results of top 1/3/5/10 molecules in drug generation.

| | Test set | GraphBP | Pocket2Mol | ResGen | AR | FLAG | TargetDiff | Decomp-o | Decomp-r | OurModel |
|---|---|---|---|---|---|---|---|---|---|---|
| **Top 1** | | | | | | | | | | |
| Vina Dock (↓) | -7.204 | -9.332 | -9.418 | -9.0326 | -8.3907 | -8.333 | -10.132 | -10.0313# | -8.387 | **-10.636** |
| Vina Score (↓) | -5.916 | -1.17 | -8.742 | -7.2451 | -8.1541 | -6.4982 | -9.8856 | -9.4223# | -7.5622 | **-10.164** |
| QED (↑) | 0.476 | 0.5559 | 0.5408 | 0.5660# | 0.5357 | **0.6191** | 0.4656 | 0.4703 | 0.5236 | 0.5696 |
| SA (↑) | 0.7275 | 0.468 | 0.74# | 0.7434 | 0.5693 | 0.6369 | 0.4979 | 0.6015 | 0.6814 | **0.7609** |
| Lipinski (↑) | 4.34 | 4.81 | 4.91# | 4.8478 | 4.6907 | **4.98** | 4.61 | 4.5253 | 4.68 | 4.94 |
| LogP | 0.8943 | 1.5523 | 2.7609 | 2.7112 | 0.6642 | 2.7022 | 2.4475 | 3.4916 | 2.1742 | 3.4859 |
| **Top 3** | | | | | | | | | | |
| Vina Dock (↓) | -7.204 | -8.809 | -9.2583 | -8.8471 | -8.19 | -8.0387 | -9.7347 | -9.6953# | -8.272 | **-10.45** |
| Vina Score (↓) | -5.916 | 2.9832 | -8.5038 | -6.9575 | -7.9136 | -6.0975 | -9.232 | -9.0352# | -7.4 | **-9.9918** |
| QED (↑) | 0.476 | 0.496 | 0.5379 | 0.5722 | 0.5251 | **0.6237** | 0.4828 | 0.4724 | 0.5252 | 0.5665# |
| SA (↑) | 0.7275 | 0.4775 | 0.7379# | 0.7554 | 0.5709 | 0.6412 | 0.5059 | 0.6 | 0.678 | **0.7684** |
| Lipinski (↑) | 4.34 | 4.787 | 4.89# | 4.8587 | 4.7285 | **4.993** | 4.5867 | 4.5657 | 4.6433 | 4.9333 |
| LogP | 0.8943 | 1.47 | 2.6559 | 2.7182 | 0.6487 | 2.4376 | 2.481 | 3.4328 | 2.073 | 3.3994 |
| **Top 5** | | | | | | | | | | |
| Vina Dock (↓) | -7.204 | -8.515 | -9.1444 | -8.7183 | -8.0713 | -7.8486 | -9.4992# | -9.5073 | -8.1604 | **-10.3422** |
| Vina Score (↓) | -5.916 | 6.5512 | -8.3641 | -6.7795 | -7.7348 | -5.8563 | -8.8716# | -8.796 | -7.2462 | **-9.8861** |
| QED (↑) | 0.476 | 0.523 | 0.5407 | 0.5706 | 0.5239 | **0.6353** | 0.4922 | 0.4708 | 0.5264 | 0.5569# |
| SA (↑) | 0.7275 | 0.478 | 0.7389# | 0.7607 | 0.5724 | 0.6432 | 0.5132 | 0.6023 | 0.6743 | **0.7798** |
| Lipinski (↑) | 4.34 | 4.776 | 4.898# | 4.8696 | 4.732 | **4.994** | 4.598 | 4.5596 | 4.628 | 4.918 |
| LogP | 0.8943 | 1.43 | 2.6084 | 2.7063 | 0.6285 | 2.3058 | 2.4093 | 3.3814 | 2.0164 | 3.3672 |
| **Top 10** | | | | | | | | | | |
| Vina Dock (↓) | -7.204 | -8.0912 | -8.9539 | -8.5095 | -7.88 | -7.5437 | -9.1306# | -9.2073 | -7.9574 | **-10.171** |
| Vina Score (↓) | -5.916 | 9.12 | -8.1363 | -6.4776 | -7.4422 | -5.4734 | -8.3162# | -8.4 | -6.9871 | **-9.7015** |
| QED (↑) | 0.476 | 0.529 | 0.5534 | 0.5707 | 0.5163 | **0.6397** | 0.4895 | 0.4725 | 0.5286 | 0.5599# |
| SA (↑) | 0.7275 | 0.485 | 0.7407# | 0.7625 | 0.5743 | 0.6442 | 0.5248 | 0.604 | 0.67 | **0.7759** |
| Lipinski (↑) | 4.34 | 4.778 | 4.901# | 4.8609 | 4.731 | **4.993** | 4.609 | 4.5182 | 4.609 | 4.919 |
| LogP | 0.8943 | 1.366 | 2.5724 | 2.5840 | 0.5792 | 2.1185 | 2.2527 | 3.2342 | 1.8905 | 3.3046 |

### 6 Detailed Interaction pattern analysis for therapeutic targets

Figure S1 demonstrates Rag2Mol's ability to capture microscopic interaction patterns on three therapeutic targets: AKT1 (PDB id: 4gv1), CDK2 (PDB id: 1h00), and AROK (PDB id: 1zyu). Rag2Mol-generated molecules display reasonable binding poses, suggesting effective inference of hit positioning within the protein pocket. Using PLIP, we analyze interactions between these targets and Rag2Mol-generated ligands, comparing them to experimentally validated active ligands. Rag2Mol reproduces most key interactions observed in experimental ligands (6/7 for 4gv1, 4/6 for 1h00, 3/6 for 1zyu) and predicts additional, physically plausible interactions (e.g., 4 and 5 extra interactions for 4gv1 and 1zyu, respectively), enhancing binding potential. Detailed interaction statistics are provided in Tables S5-S7.

For the CDK2 target, Rag2Mol preserves critical interactions, including hydrophobic contacts and hydrogen bonds with ASP145 and LEU83, while capturing an extra water-mediated bridge involving LYS20. For AROK, Rag2Mol retains all water bridges (ARG110, ARG117) and π-cation interaction (ARG110), and also favors phosphorus atom generation. While it generates five additional hydrogen bonds. Similarly, for AKT1, Rag2Mol accurately reproduced hydrophobic interactions with ALA177 and LYS179 and hydrogen bonds with ALA230, GLU234, and ASP292, while introducing extra interactions that further enhanced binding potential. These results highlight Rag2Mol's ability to learn advanced energy distributions and interaction rules.

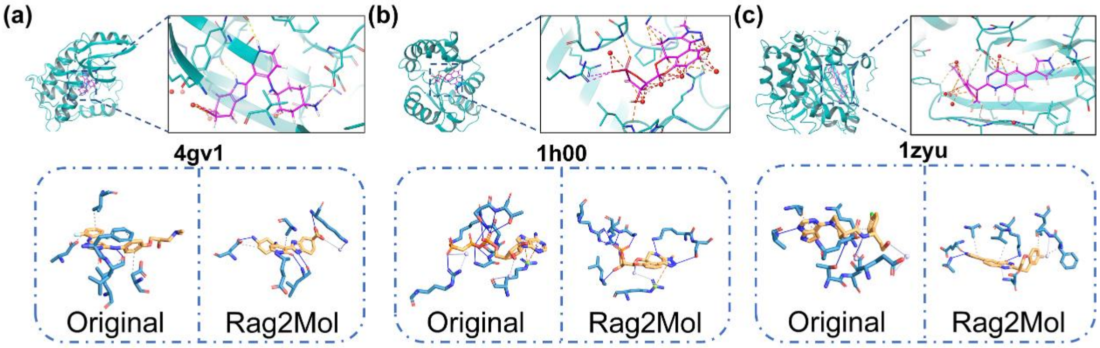


Figure S1. Protein–ligand interaction analysis. The top quadrants show the poses of Rag2Mol's generated ligands within the protein pocket, whereas the below quadrants denote the protein-ligand interaction patterns for the original and generated ligands, respectively.

#### 7 Interaction statistics

We put the detailed interaction statistics for section 3.3 and 3.5 here.

Table S5. Therapeutic target CDK2.

| 1h00 | Hydrophobic | | | | | Hydrogen Bond | | | Water Bridges |
|---|---|---|---|---|---|---|---|---|---|
| Residue | LYS20 | ALA31 | ASP86 | LEU134 | ASP145 | GLU81 | LEU83 | ASP145 | LYS20 |
| Original | √ | | √ | √ | √ | | √ | √ | |
| Rag2Mol | √ | √ | | | √ | √ | √ | √ | √ |

Table S6. Therapeutic target AROK.

| 1zyu | Hydrogen Bonds | | | | | | | | | | |
|---|---|---|---|---|---|---|---|---|---|---|---|
| Residue | SER13 | GLY14 | LYS15 | SER16 | THR17 | ARG58 | GLY80 | GLY81 | ARG117 | LEU119 | ARG136 |
| Original | √ | √ | √ | √ | √ | | | | √ | | |
| Rag2Mol | | | √ | √ | | √ | √ | √ | √ | √ | √ |

| 1zyu | Water Bridges | | **π**-Cation Interactions |
|---|---|---|---|
| Residue | ARG110 | ARG117 | ARG110 |
| Original | √ | √ | √ |
| Rag2Mol | √ | √ | √ |

Table S7. Therapeutic target AKT1.

| 4gv1 | Hydrophobic | | | | Hydrogen Bonds | | | | Water Bridges | | |
|---|---|---|---|---|---|---|---|---|---|---|---|
| Residue | PHE161 | VAL164 | ALA177 | LYS179 | GLU228 | ALA230 | GLU234 | ASP292 | GLY162 | GLU278 | ASP292 |
| Original | | | √ | √ | | √ | √ | √ | | √ | √ |
| Rag2Mol | √ | √ | √ | √ | √ | √ | √ | √ | √ | | √ |

Table S8. Real-world application PTPN2.

| 7uad | Hydrophobic | | | | | Pi-stacking | Water-bridge |
|---|---|---|---|---|---|---|---|
| Residue | TYR48 | VAL51 | PHE183 | ALA218 | ILE220 | TYR48 | ASP50 |
| Original | √ | √ | √ | √ | √ | | |
| Find: 1 | √ | √ | √ | √ | √ | √ | |
| Gen: 1 | | √ | √ | | | √ | √ |
| | Hydrogen Bonds | | | | | | |
| Residue | TYR48 | ASP50 | LYS122 | ASP182 | PHE183 | SER217 | ALA218 |
| Original | | √ | √ | √ | √ | √ | √ |
| Find: 1 | | | √ | √ | √ | √ | √ |
| Gen: 1 | √ | | √ | √ | √ | √ | √ |
| Residue | GLY219 | ILE220 | GLY221 | ARG222 | GLN264 | | |
| Original | √ | √ | √ | √ | √ | | |
| Find: 1 | √ | | | √ | √ | | |
| Gen: 1 | √ | | √ | √ | √ | | |

### 8 Connection between Rag2Mol-G and Rag2Mol-R

To demonstrate that the molecules retrieved by Rag2Mol-R inherit the favorable properties of the molecules designed by Rag2Mol-G, we compared the RMSD of the molecules obtained from both methods. As shown in Figure S2, we used RDKit to compare the differences in binding poses between AI-generated molecules and those obtained through virtual screening within each protein pocket in the test set. A total of 17,470 pairs of generated and screened molecules were analyzed across 100 target protein pockets, with an average RMSD of 0.7217 and a maximum RMSD of 3.7881.

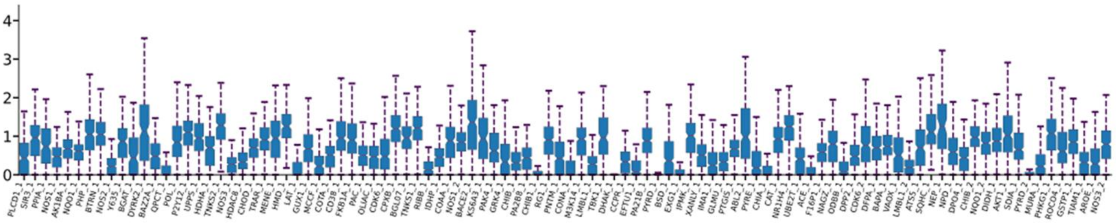


Figure S2. The RMSD comparison between Rag2Mol-G and Rag2Mol-R

### 9 Connection between retrieved and generated molecules

We design experiments to demonstrate that Rag2Mol learns hidden topological knowledge and interaction patterns from reference molecules rather than merely reusing structures. As shown in Figure S3, we analyze the similarity between the retrieved reference molecules and all baseline-generated molecules. Compared to other AI methods without reference molecules, Rag2Mol-generated molecules did not exhibit outstanding similarity performance.

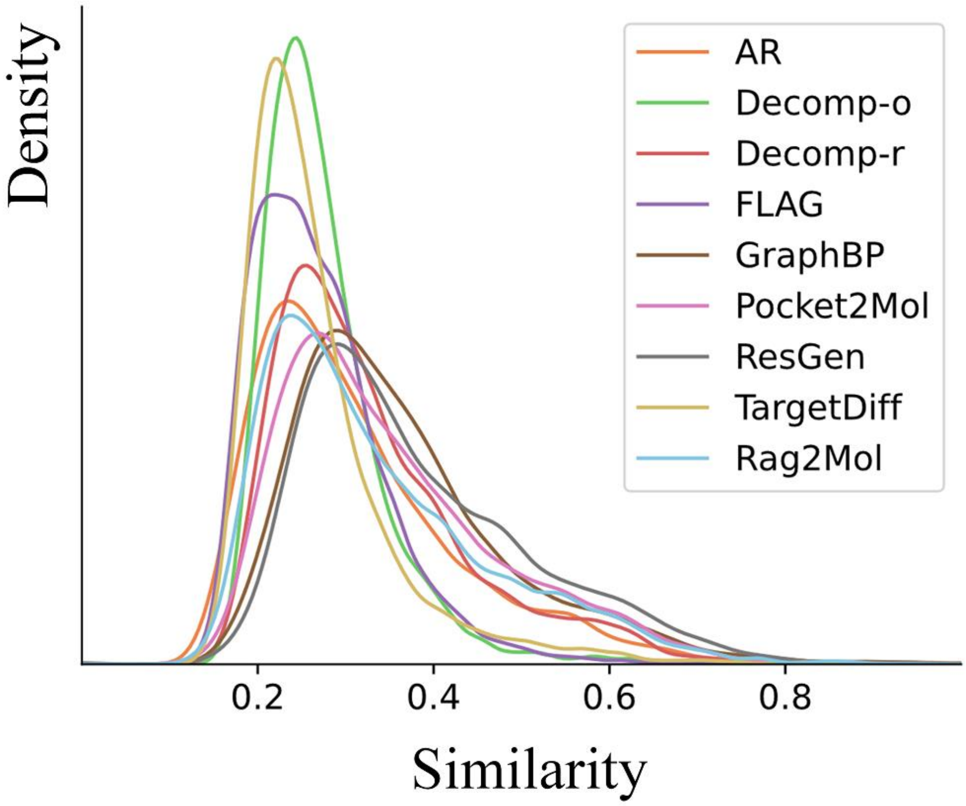


Figure S3. The similarity between the retrieved reference molecules and AI-generated molecules.

### 10 Impact of different databases

We re-apply Rag2Mol-G on test set by feeding different retrieved molecular databases. Table S9 shows little difference in the generated results based on different databases. This is due to the high similarity of molecules retrieved in different databases. Table S10 shows the molecular similarity retrieved from different databases.

Table S9. The top 10 molecules mean binding energies and drug-likeness properties

| | Test set | ChEMBL | GEOM | ZINC15 | BindingDB |
|---|---|---|---|---|---|
| Affinity (↑) | - | 60.29 % | 60.78 % | 60.99 % | **61.52 %** |
| Vina Dock (↓) | -7.204 | -9.9918 | -10.0052 | -9.9829 | **-10.171** |
| QED (↑) | 0.476 | 0.5684 | 0.4725 | **0.5737** | 0.5599 |
| SA (↑) | 0.7275 | **0.8068** | 0.7966 | 0.7941 | 0.7759 |
| Lipinski (↑) | 4.34 | **4.94** | 4.927 | 4.926 | 4.919 |
| LogP | 0.8943 | 3.3148 | 3.2032 | 3.1931 | 3.3046 |

Table S10. Molecular similarity retrieved from different databases.

| | ChEMBL | GEOM | ZINC15 | BindingDB |
|---|---|---|---|---|
| ChEMBL | 1.0 | 0.7064 | 0.6029 | 0.629 |
| GEOM | 0.6121 | 1.0 | 0.5929 | 0.6 |
| ZINC15 | 0.5736 | 0.6407 | 1.0 | 0.5359 |
| BindingDB | 0.6144 | 0.6597 | 0.5476 | 1.0 |

To further elucidate the influence of different databases, distinct databases are utilized for the purposes of RAG and subsequent searches. For convenience, only the average Vina docking scores and QED values of the top 10 molecules in each database pair are provided here. In Table S11 and S12, the RAG databases are arranged in rows and the similarity search databases are arranged in columns.

Table S11. Vina Dock score for the top 10 molecules for each pair of databases.

| | ChEMBL | GEOM | ZINC15 | BindingDB |
|---|---|---|---|---|
| ChEMBL | -9.6863 | -9.7091 | -9.6603 | -9.9088 |
| GEOM | -9.5638 | -9.5914 | -9.5757 | -9.8 |
| ZINC15 | -9.7693 | -9.7969 | -9.7329 | -9.939 |
| BindingDB | -9.8207 | -9.8681 | -9.84 | -10.0471 |

Table S12. QED score for the top 10 molecules for each pair of databases.

| | ChEMBL | GEOM | ZINC15 | BindingDB |
|---|---|---|---|---|
| ChEMBL | 0.6442 | 0.6439 | 0.6474 | 0.6378 |
| GEOM | 0.6648 | 0.6629 | 0.6703 | 0.6537 |
| ZINC15 | 0.697 | 0.7049 | 0.7014 | 0.6909 |
| BindingDB | 0.5366 | 0.5443 | 0.5453 | 0.5071 |
